# Number of days required to estimate objectively measured physical activity constructs in different age groups

**DOI:** 10.1101/610030

**Authors:** Luiza Isnardi Cardoso Ricardo, Andrea Wendt, Leony Morgana Galliano, Werner de Andrade Muller, Gloria Izabel Niño Cruz, Fernando Wehrmeister, Soren Brage, Ulf Ekelund, Inácio Crochemore M Silva

## Abstract

The present study aims to estimate the minimum number of accelerometer measurement days needed to estimate habitual physical activity (PA) among 6, 18 and 30 years old participants, belonging to three population-based Brazilian birth cohorts. Method: PA was assessed by triaxial wrist worn GENEActiv accelerometers for 4-7 consecutive days, including at least one weekend day. Accelerometer raw data were analyzed with R-package GGIR. Description of PA measures (overall PA, MVPA, LPA) between weekdays and weekend days were conducted, and statistical differences were tested with chi-squared and Kruskal-Wallis tests. Intraclass Reliability Coefficient (IRC) was applied through the Spearman Brown Formulae to test reliability of different number of days of accelerometer use. Results: Differences between week and weekend days regarding LPA, MVPA and Overall PA, were only observed among 30-year-olds. More MVPA (p=0.006) and Overall PA (p<0.001) were performed on week days. The IRC coefficients ranged from 0.44 to 0.83 in children and from 0.54 to 0.88 in adults. Conclusion: In conclusion, our results show that between four and six measurement days are needed to achieve good reliability in the analyzed participants, depending on the PA construct evaluated (MVPA, LPA or overall PA).

## Introduction

Physical activity (PA) has positive effect on health and quality of life of individuals and communities ^1^. Evidence shows that PA is a protective behavior on major non-communicable diseases, such as coronary heart disease, type II diabetes and cancer ^2^. However, global PA prevalence is still low ^3^, including such behavior in the public health agenda and proving the need of monitoring PA at population levels.

Accelerometers (portable motion sensors) have been increasingly adopted in large scale studies, since it provides more accurate physiological and mechanical parameters to estimate PA ^4^. In the last decade, the use of accelerometers for this purpose increased significantly ^5^, due to the capability to quantify duration, frequency and intensity of PA through acceleration signals, movement patterns and its magnitude ^6^. Also, currently available accelerometers enables large quantity of data storage, a wide spectrum of cut-off points for different PA intensities, movement pattern recognition and the possibility of more detailed analyzes using the raw data ^5^.

In order to better understand the prevalence, levels and impact of PA on health, accurate measures are essential. However, with the increase of accelerometer-based research, the variability among protocols is also rising, since there is no standardized recommendation for data collection. There are different sources of variability, such as accelerometers brands, placements, unit of analyzes, minimum number of hours per day and minimum number of measurement days. All these decisions will influence final PA estimates at some degree, and therefore must be discussed accordingly to their advantages and disadvantages.

One of these methodological decisions is the minimum number of days of measurement, which is an important decision that will define the logistics of each study and the representativeness of habitual PA behavior. However, it seems that there is no consensus on the literature. Among adults, at least two days for light and moderate PA ^7^and three days for moderate-to-vigorous PA seems to be needed ^8^. Additionally, Scheers et al.^9^ suggests that PA should be measured for two weekend days and three weekdays in order to achieve common week reliability in adults. Among children, the widely accepted recommendation seems to be between four and five measurement days ^10^.

Historically, considering that during the first two decades of accelerometer use these measurements have been conducted by waist/hip-worn devices, decision regarding the minimum number of days to estimate habitual physical activity was usually associated with a minimum number of hours per day criteria. However, studies using wrist-worn accelerometers usually adopt a 24 hours-protocol with increased compliance (Troiano et al., 2014). Therefore, the present study aims to estimate the minimum number of accelerometer measurement days needed to estimate habitual PA among 6, 18 and 30 years old participants, belonging to three population-based Brazilian birth cohorts.

## Materials and Methods

### Study design and participants

This study is based on data from three birth cohorts studies conducted in Pelotas (a southern city in Brazil) where PA was objectively assessed between 2010 and 2012. All babies delivered in hospital in 1982, 1993 and 2004 were identified during daily visits to the maternity hospitals in the city (more than 99% of deliveries occur in hospitals). In each cohort, less than 1% of participants recruited refused to participate. All three cohorts have been followed up at different time points thereafter. More details regarding the methodology of each cohort have been previously described elsewhere ^11–13^.

Recent follow-up visits were performed when cohort participants born in 1982, 1993 and 2004 were approximately 30 years, 18 years and 6 years old, respectively. At these follow-ups, participants were interviewed and clinically examined by the research team. Data on socioeconomic and demographic characteristics, anthropometry, clinical and biochemical measurements were collected. After the clinical examinations, participants were invited to wear an accelerometer (GENEActiv; ActivInsights, Kimbolton, UK) on the non-dominant wrist.

Approval for the study was obtained from the Institutional Ethics Committee and all participants or their legal representatives voluntarily signed written informed consent.

### Physical activity measurement

PA was assessed by triaxial wrist worn accelerometers, allowing measures of body movements on three axes: vertical (Y), horizontal right-left (X), and horizontal front-back axis (Z), within an acceleration dynamic range of ± 8g ^14^, where g represents gravitational unit. Sampling frequency was set at 85.7 Hz.

For practical reasons and to increase compliance, participants were invited to wear the accelerometer using a 24 hours protocol (during awakening and sleeping hours) for 4-7 consecutive days, including at least one weekend day. The total amount of monitored days varied according to the day of the clinical visit. Participants who visited the clinic on Mondays, Tuesdays or Wednesdays were monitored until the following Monday, whereas those who visited the clinic on Thursdays, Fridays or Saturdays, were monitored until the following Wednesday. Further information regarding the protocol is available in a previous publication ^15^. In the present study, all analyzes were restricted to individuals who provided six full days of data.

### Data reduction

Accelerometers were set up and downloaded in the GENEActiv software. Accelerometer data in binary format were analyzed with R-package GGIR [http:/cran.r-project.org] ^16^. The detailed signal processing scheme included the following steps: verification of sensor calibration error using local gravity as a reference; detection of sustained abnormally high values and non-wear detection. Furthermore, calculation of the vector magnitude of activity-related acceleration Euclidian Norm Minus One (ENMO) was used to summarize three-dimensional raw data (from axes x, y, and z) into a single-dimensional signal vector magnitude (ENMO = ∑ | – 1g|). The data were further summarized when calculating the average values per 5-second-epochs. The summary measures used were (1) the average ENMO per day (expressed in mg and as an estimate of overall PA volume), (2) estimated time spent in 10-minute-bouted moderate-to-vigorous PA (MVPA) per day and, (3) time spent in non-bouted light PA (LPA) (expressed in minutes). MVPA was defined as ENMO records above 100mg, while bouts criterion was defined as consecutive periods in which participants spent at least 80% of time in MVPA. LPA was considered activities with acceleration between 50mg and 100mg.

### Statistical analyzes

Descriptive analyzes comparing characteristics of individuals with six complete days of data and the remaining participants of the cohorts were performed using ANOVA or its non-parametric equivalent when necessary, presenting absolute and relative frequencies distribution and respective p values, according to sex (male/female), weight status (underweight, normal-weight, overweight and obese) based on the World Health Organization ^17^ classification of BMI for adults (1982 cohort) and according to the age and sex Z-scores for BMI for children and adolescents (1993 and 2004 cohorts) ^18^.

Also, description of PA measures (overall PA, MVPA, LPA) between weekdays and weekend days were conducted, and statistical differences were tested with chi-squared and Kruskal-Wallis tests. Intraclass Reliability Coefficient (IRC) was applied through the Spearman Brown Formulae ^19^to test reliability of different number of days of accelerometer use. All analyzes were stratified by age groups (birth cohorts) and performed in the software Stata 12.0. Statistical significance was set at 5%.

## Results

Sample description, as well as comparison between individuals with six measurement days (analytical sample) and the remaining members followed-up on the most recent data collection of each cohort is presented in Table 1. There were no significant differences between the analytical sample and the rest of the cohort for most variables, except for gender among 18-year-olds whose presented a higher proportion of males (54.9).

**Table 1.** Sample characteristics of six, 18 and 30-year-old individuals that wore the accelerometer for six and less than six measurement days

| Variables | 6 years (N=103) |  |  | 18 years (N=504) |  |  | 30 years (N=453) |  |  |
| --- | --- | --- | --- | --- | --- | --- | --- | --- | --- |
|  | < 6 days | 6 days | p | < 6 days | 6 days | P | < 6 days | 6 days | p |
| **Gender** |  |  | 0.132 |  |  | 0.005 |  |  | 0.410 |
| Male | 1,294 (51.8) | 47 (45.6) |  | 1,487 (48.1) | 276 (54.9) |  | 1,099 (48.6) | 210 (46.5) |  |
| Female | 1,206 (48.2) | 56 (54.4) |  | 1,603 (51.9) | 227 (45.1) |  | 1,161 (51.4) | 242 (53.5) |  |
| **Asset index**<br>**(quintiles)** |  |  | 0.147 |  |  | 0.741 |  |  | 0.349 |
| 1° (poorest) | 246 (10.1) | 10 (9.8) |  | 616 (20.1) | 104 (20.7) |  | 514 (24.4) | 121 (28.7) |  |
| 2° | 371 (15.3) | 23 (22.6) |  | 615 (20.0) | 91 (18.1) |  | 453 (21.5) | 86 (20.4) |  |
| 3° | 574 (23.6) | 15 (14.7) |  | 608 (19.8) | 109 (21.7) |  | 566 (26.8) | 100 (23.7) |  |
| 4° | 489 (20.1) | 22 (21.6) |  | 631 (20.5) | 105 (20.9) |  | 195 (9.2) | 42 (10.0) |  |
| 5° (richest) | 750 (30.9) | 32 (31.4) |  | 603 (19.6) | 93 (18.5) |  | 383 (18.1) | 73 (17.3) |  |
| **BMI** |  |  | 0.060 |  |  | 0.600 |  |  | 0.958 |
| Normal | 1,487 (63.9) | 70 (72.9) |  | 2,189 (72.3) | 368 (73.9) |  | 889 (41.0) | 175 (40.3) |  |
| Overweight | 434 (18.7) | 18 (18.8) |  | 543 (17.9) | 80 (16.1) |  | 755 (34.8) | 154 (35.5) |  |
| Obesity | 405 (17.4) | 8 (8.3) |  | 297 (9.8) | 50 (10.0) |  | 524 (24.2) | 105 (24.2) |  |

Figure 1 presents the mean and median of minutes spent per day in LPA and MVPA, respectively, and the mean overall PA. Among adults (18 and 30 years old), overall PA, MVPA and LPA median were lower on Sundays compared to the rest of the week. Among children, MVPA was lowest on Mondays. Differences between week and weekend days regarding LPA, MVPA and Overall PA, were only observed among 30-year-olds. More MVPA (p=0.006) and Overall PA (p<0.001) were performed on week days (Figure 2).

**Figure 1.**
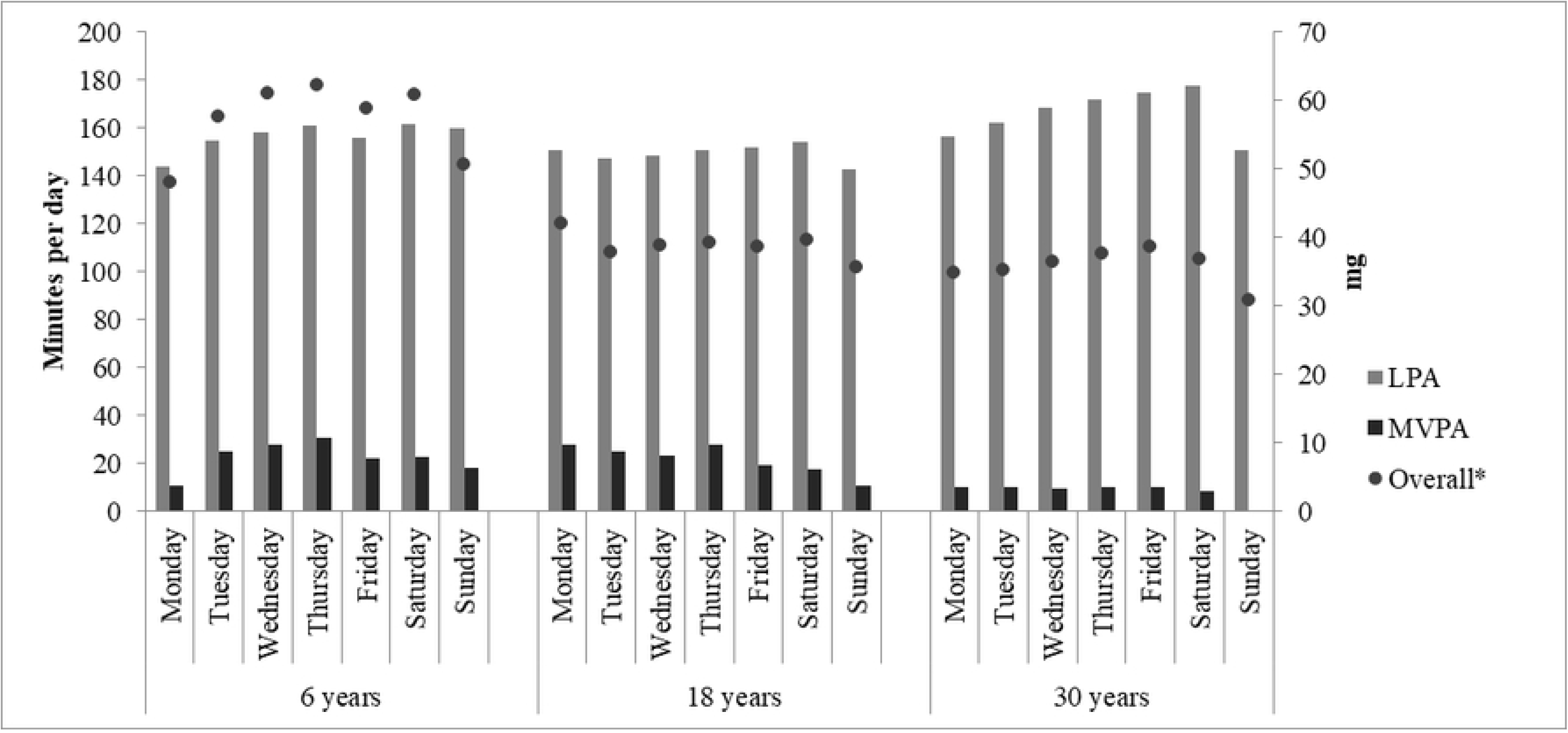

**Figure 2.**
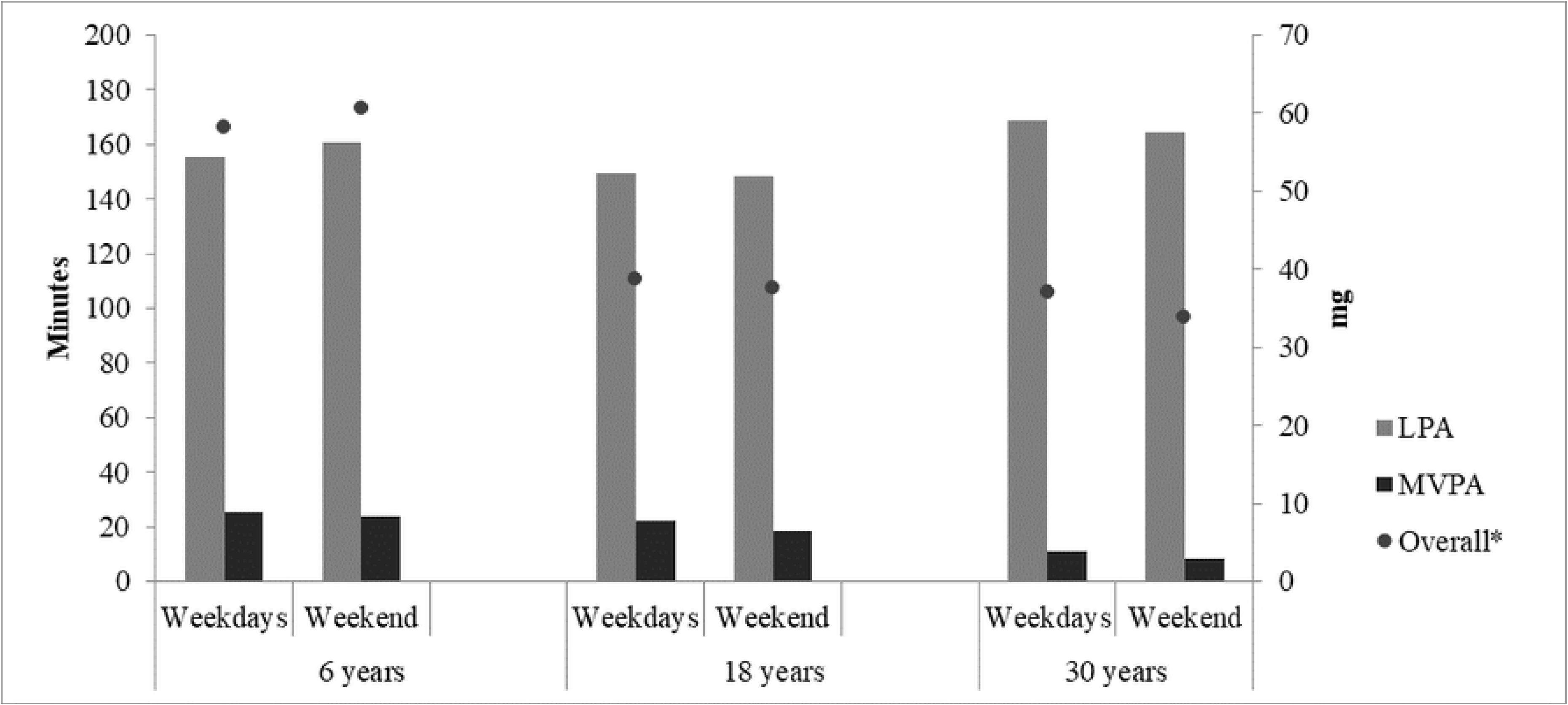

Figures 3A, 3B and 3C show the Intraclass Reliability Coefficient (IRC) for one to six measurement days. Highest IRC coefficients was observed for overall PA in all three groups. The coefficients ranged from 0.44 to 0.83 in children and from 0.54 to 0.88 in adults. To achieve an IRC >0,7 two and three days of measurements were needed in adults and children, respectively. Three and four days of measurements of MVPA was needed to achieve an IRC >0,7 in 30-year-olds and in 18 and 6-year-olds, respectively. Five days of LPA was necessary to reach an IRC>0.7 in 6-year-old children, three days for 18 years-old adults, and four days for 30 years-old adults. To achieve an IRC>0.8, four and five days of overall PA was needed among adults and children, respectively.

**Figure 3.**
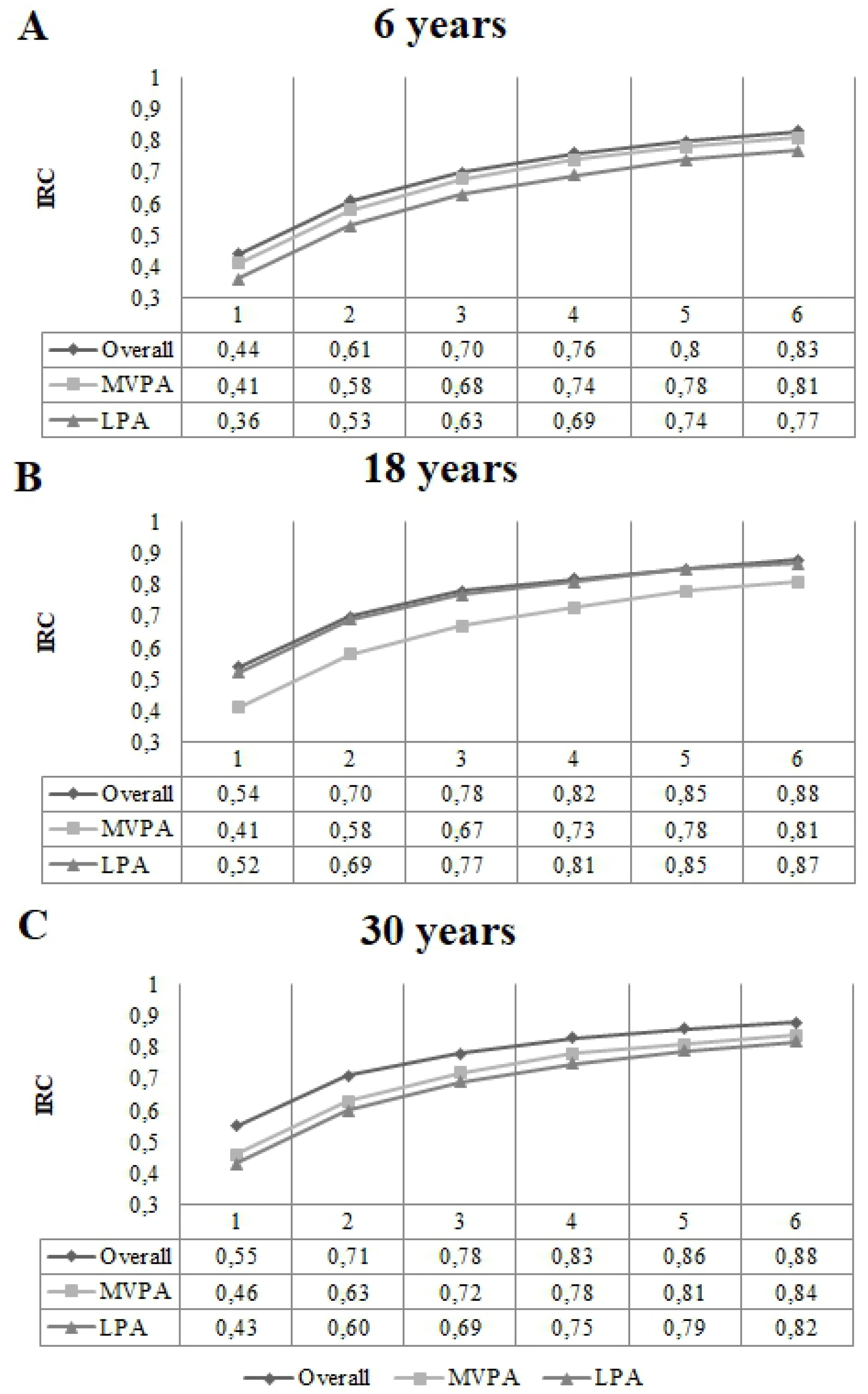

## Discussion

Our results show that, for MVPA, four days would be necessary to represent a week of measurement using the 0.7 threshold; and 5 or 6 days would be needed using the 0.8 threshold. These results vary according to the construct evaluated (MVPA, LPA or overall PA). Overall PA follows a stable pattern through the week, resulting in a smaller number of days needed to an admissible IRC.

A better interpretation of our results requires the understanding of the main differences among the PA constructs analyzed. For instance, overall PA demonstrates the behavior with minimum arbitrariness, including all daily movement, including both daily life activities and a more structured PA. Similarly, light PA as treated in the present study, non-bouted, represents low intensity activities without any duration restriction, but excluding sedentary time and MVPA. Lastly, when using 10 minutes bouts, MVPA estimates attempt to represent structured activities performed for longer periods. Therefore, it is expected to find different reliability results among the outcomes, once they represent distinct constructs of PA.

We observed no differences in any of the PA constructs between weekdays and weekend days, except for 30-year-old adults, in which MVPA and overall PA levels were lower on weekend days. Likewise, adults presented lower PA on Sundays in the daily comparison analyzes. This may occur due to the decrease in workload and routine activities, since most people work from Monday to Saturday. For 18 years old adults, the analyzes between weekend and weekdays’ PA did not present significant difference, possibly because Saturday presents similar results than the remaining week, diluting the difference between weekdays and weekend days. Also, children may present different patterns of PA then adults when comparing weekdays and weekend days, since during weekdays there is more participation in school activities, which may impact positively on children’s PA ^20^. Opposite to our results, other studies have reported differences regarding weekdays and weekend days among children and adults ^21–23^. This may be due to our specific sample, which includes only 6, 18 and 30-year-old individual, not an age range (e.g. 18 to 60-year-old individuals). However, it is important to highlight that reliability is not just an age-related phenomenon, it is also a cultural aspect and observed reliability estimates in the present study does not necessarily translate to other settings.

Most previous studies examining reliability, or other measures to establish the minimum number of measurement days to achieve a desirable IRC have usually examined different PA intensities such as light, moderate, vigorous or MVPA ^9,21^. However, it is also important to assess overall PA, which integrates information regarding all intensities with lower arbitrariness, presenting good metrics of different intensity categories, since it incorporates a continuum of PA intensities performed ^8^. Furthermore, overall PA expressed as mg may improve a direct comparison between studies ^24^.

Regarding children, more days of measurement were necessary to achieve IRC>0.7, for all three PA measures. This could be due to higher variability of our children sample compared to adults and other studies with children. Addy et.al. ^25^ found that 3.6 days of measurement would be needed to achieve an ICC of 0.7 and 6.2 would be necessary for an ICC of 0.8 among preschool children. Another study, using wrist-worn accelerometers showed an IRC of 0.8 with two days evaluating adults’ MVPA ^21^. This evidence shows that many factors could impact on the decision of how many days should be included in accelerometry data analyzes, such as: a) IRC ideal threshold, b) study population, c) accelerometer model and placement, and d) the PA construct of interest. Therefore, it is important to consider an accelerometry protocol including as many measurement days as possible, in order to guarantee good reliability. Also, different days of protocol are needed depending on the research question. Surveillance studies may be satisfied with only one day of data while studies about seasonality need many days in different periods of year. Furthermore, the present analyzes are recommended for each data collection *a posteriori*, in order to define the minimum of valid days of measurement in each study.

The present study, based on large birth cohorts born 11 years apart, with high follow-up rates and objective measures of PA, provides reliable estimates of the number of days needed to obtain desirable IRC. However, some limitations need to be addressed, such as the decrease in the number of observations when restricting the analyzes to those with six consecutive days, mainly among children. This decrease is due to the use of the non-commercial version of the GENEA accelerometer which is not comparable to the GENEActiv and were not analyzed in the present study.

In conclusion, our results show that between four and six measurement days are needed to achieve good reliability in the analyzed participants, depending on the PA construct evaluated (MVPA, LPA or overall PA). However, final decisions on the research protocol of accelerometer data collection should balance according to each research question, available logistic and potential losses of valid days during the process.

### What does this article add?

Since we found differences between weekdays and weekend days physical activity among adults, it is important to make efforts towards weekend days measurement in order to well represent the physical activity variability across the week. The key finding is that the number of measurement days required to estimate physical activity depends on the age range and the construct of physical activity evaluated, varying between four and six measurement days. When evaluating children’s physical activity, there is a need for more days of measurement when compared to adults, regardless of the construct of physical activity. From a practical point of view, we recommend measuring as many days as possible to guarantee good reliability.

## Acknowledgements

This article is based on data from the studies “Pelotas Birth Cohort, 1982”, “Pelotas Birth Cohort, 1993” and “Pelotas Birth Cohort, 2004” conducted by Postgraduate Program in Epidemiology at Universidade Federal de Pelotas with the collaboration of the Brazilian Public Health Association (ABRASCO). From 2004 to 2013, the Wellcome Trust supported all three birth cohort studies. National Support Program for Centers of Excellence (PRONEX), the Brazilian National Research Council (CNPq), and the Brazilian Ministry of Health supported previous phases of the studies. The 1982 Birth Cohort were funded by the International Development Research Center, Overseas Development Administration. The World Health Organization funded the 1982 and 2004 Birth cohorts. And the 1982 and 1993 Birth Cohort were also supported by the European Union.

## References

1. Arena R, McNeil A, Sagner M, Hills AP. The Current Global State of Key Lifestyle Characteristics: Health and Economic Implications. Prog Cardiovasc Dis [Internet]. 2017 Mar [cited 2019 Apr 10];59(5):422–9. Available from: https://linkinghub.elsevier.com/retrieve/pii/S0033062017300257

2. Lee I-M, Shiroma EJ, Lobelo F, Puska P, Blair SN, Katzmarzyk PT. Effect of physical inactivity on major non-communicable diseases worldwide: an analysis of burden of disease and life expectancy. Lancet [Internet]. 2012 Jul [cited 2019 Apr 10];380(9838):219–29. Available from: https://linkinghub.elsevier.com/retrieve/pii/S0140673612610319

3. Sallis JF, Bull F, Guthold R, Heath GW, Inoue S, Kelly P, et al. Progress in physical activity over the Olympic quadrennium. Lancet. 2016;388(10051):1325– 36.

4. Ainsworth B, Cahalin L, Buman M, Ross R. The Current State of Physical Activity Assessment Tools. Prog Cardiovasc Dis [Internet]. 2015 Jan [cited 2019 Apr 10];57(4):387–95. Available from: https://linkinghub.elsevier.com/retrieve/pii/S0033062014001674

5. Troiano RP, McClain JJ, Brychta RJ, Chen KY. Evolution of accelerometer methods for physical activity research. Br J Sports Med [Internet]. 2014 Jul [cited 2018 Oct 2];48(13):1019–23. Available from: http://www.ncbi.nlm.nih.gov/pubmed/24782483

6. Chen KY, Bassett DR. The technology of accelerometry-based activity monitors: current and future. Med Sci Sports Exerc [Internet]. 2005 Nov [cited 2018 Nov 26];37(11 Suppl):S490–500. Available from: http://www.ncbi.nlm.nih.gov/pubmed/16294112

7. Dillon CB, Fitzgerald AP, Kearney PM, Perry IJ, Rennie KL, Kozarski R, et al. Number of Days Required to Estimate Habitual Activity Using Wrist-Worn GENEActiv Accelerometer: A Cross-Sectional Study. Buchowski M, editor. PLoS One [Internet]. 2016 May 5 [cited 2019 Apr 10];11(5):e0109913. Available from: https://dx.plos.org/10.1371/journal.pone.0109913

8. Matthews CE, Keadle SK, Troiano RP, Kahle L, Koster A, Brychta R, et al. Accelerometer-measured dose-response for physical activity, sedentary time, and mortality in US adults. Am J Clin Nutr [Internet]. 2016 Nov 1 [cited 2019 Apr 10];104(5):1424–32. Available from: https://academic.oup.com/ajcn/article/104/5/1424/4564391

9. Scheers T, Philippaerts R, Lefevre J. Variability in physical activity patterns as measured by the SenseWear Armband: how many days are needed? Eur J Appl Physiol [Internet]. 2012 May 28 [cited 2019 Apr 10];112(5):1653–62. Available from: http://link.springer.com/10.1007/s00421-011-2131-9

10. Kang M, Bjornson K, Barreira T V, Ragan BG, Song K. The minimum number of days required to establish reliable physical activity estimates in children aged 2– 15 years. Physiol Meas [Internet]. 2014 Nov 1 [cited 2019 Apr 10];35(11):2229–37. Available from: http://stacks.iop.org/0967-3334/35/i=11/a=2229?key=crossref.5f1de6ba6f0519a711abeee812a5bfd6

11. Gonçalves H, Assunção MC, Wehrmeister FC, Oliveira IO, Barros FC, Victora CG, et al. Cohort Profile update: The 1993 Pelotas (Brazil) Birth Cohort followup visits in adolescence. Int J Epidemiol [Internet]. 2014 Aug [cited 2018 Dec 12];43(4):1082–8. Available from: http://www.ncbi.nlm.nih.gov/pubmed/24729426

12. Horta BL, Gigante DP, Goncalves H, dos Santos Motta J, Loret de Mola C, Oliveira IO, et al. Cohort Profile Update: The 1982 Pelotas (Brazil) Birth Cohort Study. Int J Epidemiol [Internet]. 2015 Apr 1 [cited 2019 Apr 10];44(2):441–441e. Available from: https://academic.oup.com/ije/article-lookup/doi/10.1093/ije/dyv017

13. Santos IS, Barros AJ, Matijasevich A, Zanini R, Chrestani Cesar MA, Camargo-Figuera FA, et al. Cohort Profile Update: 2004 Pelotas (Brazil) Birth Cohort Study. Body composition, mental health and genetic assessment at the 6 years follow-up. Int J Epidemiol [Internet]. 2014 Oct 1 [cited 2018 Dec 12];43(5):1437–1437f. Available from: http://www.ncbi.nlm.nih.gov/pubmed/25063002

14. Jefferis BJ, Parsons TJ, Sartini C, Ash S, Lennon LT, Wannamethee SG, et al. Does duration of physical activity bouts matter for adiposity and metabolic syndrome? A cross-sectional study of older British men. Int J Behav Nutr Phys Act [Internet]. 2016 Dec 15 [cited 2019 Apr 10];13(1):36. Available from: http://www.ijbnpa.org/content/13/1/36

15. da Silva IC, van Hees VT, Ramires V V, Knuth AG, Bielemann RM, Ekelund U, et al. Physical activity levels in three Brazilian birth cohorts as assessed with raw triaxial wrist accelerometry. Int J Epidemiol [Internet]. 2014 Dec [cited 2019 Apr 10];43(6):1959–68. Available from: https://academic.oup.com/ije/article-lookup/doi/10.1093/ije/dyu203

16. van Hees VT, Gorzelniak L, Dean León EC, Eder M, Pias M, Taherian S, et al. Separating Movement and Gravity Components in an Acceleration Signal and Implications for the Assessment of Human Daily Physical Activity. Müller M, editor. PLoS One [Internet]. 2013 Apr 23 [cited 2018 Nov 26];8(4):e61691. Available from: https://dx.plos.org/10.1371/journal.pone.0061691

17. WHO. (2000). Obesity: preventing and managing the global epidemic: World Health Organization. - Pesquisa Google [Internet]. 2000 [cited 2019 Apr 10]. Available from: https://www.google.com/search?rlz=1C1SQJL_pt-BRBR799BR799&ei=Lt-tXKaTIfa75OUP5vyqwA4&q=WHO.+%282000%29.+Obesity%3A+preventing_+and+managing+the+global+epidemic%3A+World+Health+Organization.&oq=WHO.+%282000%29.+Obesity%3A+preventing+and+managing+the+global+

18. Ede Onis M. Development of a WHO growth reference for school-aged children and adolescents. Bull World Health Organ [Internet]. 2007 Sep 1 [cited 2019 Apr 10];85(09):660–7. Available from: http://www.who.int/bulletin/volumes/85/9/07-043497.pdf

19. Eisinga R, Grotenhuis M te, Pelzer B. The reliability of a two-item scale: Pearson, Cronbach, or Spearman-Brown? Int J Public Health [Internet]. 2013 Aug 23 [cited 2019 Apr 10];58(4):637–42. Available from: http://link.springer.com/10.1007/s00038-012-0416-3

20. Berglind D, Tynelius P. Objectively measured physical activity patterns, sedentary time and parent-reported screen-time across the day in four-year-old Swedish children. BMC Public Health [Internet]. 2018 Dec 1 [cited 2019 Apr 10];18(1):69. Available from: http://bmcpublichealth.biomedcentral.com/articles/10.1186/s12889-017-4600-5

21. Dillon CB, Fitzgerald AP, Kearney PM, Perry IJ, Rennie KL, Kozarski R, et al. Number of Days Required to Estimate Habitual Activity Using Wrist-Worn GENEActiv Accelerometer: A Cross-Sectional Study. Buchowski M, editor. PLoS One [Internet]. 2016 May 5 [cited 2018 Oct 9];11(5):e0109913. Available from: http://dx.plos.org/10.1371/journal.pone.0109913

22. Doherty A, Jackson D, Hammerla N, Plötz T, Olivier P, Granat MH, et al. Large Scale Population Assessment of Physical Activity Using Wrist Worn Accelerometers: The UK Biobank Study. Buchowski M, editor. PLoS One [Internet]. 2017 Feb 1 [cited 2019 Apr 10];12(2):e0169649. Available from: https://dx.plos.org/10.1371/journal.pone.0169649

23. Nilsson A, Anderssen SA, Andersen LB, Froberg K, Riddoch C, Sardinha LB, et al. Between- and within-day variability in physical activity and inactivity in 9- and 15-year-old European children. Scand J Med Sci Sports [Internet]. 2008 Feb 3 [cited 2019 Apr 10];19(1):10–8. Available from: http://doi.wiley.com/10.1111/j.1600-0838.2007.00762.x

24. Bassett DR, Troiano RP, Mcclain JJ, Wolff DL. Accelerometer-based Physical Activity. Med Sci Sport Exerc [Internet]. 2015 Apr [cited 2019 Apr 10];47(4):833–8. Available from: https://insights.ovid.com/crossref?an=00005768-201504000-00020

25. Addy CL, Trilk JL, Dowda M, Byun W, Pate RR. Assessing Preschool Children’s Physical Activity: How Many Days of Accelerometry Measurement. Pediatr Exerc Sci [Internet]. 2014 Feb [cited 2018 Oct 9];26(1):103–9. Available from: http://www.ncbi.nlm.nih.gov/pubmed/24092773

